# The ability of enzymes to preserve reactive conformations with well-positioned residues during enzyme-catalyzed reactions can be an important factor for efficient catalysis: a case study of CARNMT1

**DOI:** 10.1101/2024.07.18.604158

**Authors:** Mingling Yang, Hao Deng, Adua Rahman, Ping Qian, Hong Guo

## Abstract

Determining the origin of high catalytic efficiency of enzymes has been a central goal in biochemical research. But our understanding of the mechanisms by which enzymes achieve enormous rate enhancements is still incomplete. One of wildly accepted strategies to achieve rate enhancements by enzymes is to position catalytic residues precisely in enzyme-substrate complexes and form reactive conformations for catalysis. Such catalytic strategy through positioning may also serve as the basis for transition state stabilization. Although considerable efforts have been made to determine relationships between enzyme activities and the formation of reactive conformations of enzyme-substrate complexes, there is considerable uncertainty concerning the ability of enzymes to preserve reactive conformations during enzyme-catalyzed reactions. The failure of properly maintaining reactive conformations during reactions may prevent residues from exerting catalytic effects and lead to significant reductions of activities. Here QM/MM MD and free energy simulations are performed for CARNMT1 to determine the importance and requirements for preservation of catalytic competent reactive conformations during enzyme-catalyzed reactions. For CARNMT1, the existence of wild-type reactive conformation is directly related to the conformation of His347. Therefore, the conformation of His347 can be used to monitor preservation or collapse of the reactive conformation during catalysis. It is found that for wild-type the reactive conformation can not only be formed in the enzyme-substrate complex but also be preserved well after the reaction passes TS. For the Tyr386Ala, Tyr396Asp and Tyr398Ala mutants, the wild-type like reactive conformations can be formed in the mutant-substrate complexes but collapse in the early stage of reaction. Such pre-mature collapse of the reactive conformation in the mutants prevents the residues to fully exert catalytic effects and leads to a significant reduction of catalytic efficiency. The results suggest that the ability of enzymes to preserve reactive conformations with well-positioned catalytic residues during enzyme-catalyzed reactions can be an important factor for efficient catalysis, and this property of enzymes deserves attention.

## Introduction

Understanding how enzymes achieve enormous rate enhancements in catalyzing chemical reactions is of fundamental importance, and a variety of potential strategies used by enzymes have been proposed to explain enzyme’s catalytic efficiencies.(1, 2) Nevertheless, our understanding of the origin of enzyme’s catalytic powers is still incomplete, and the lack of such knowledge may have contributed to the difficulty in designing highly active enzymes with efficiencies that rival the rate enhancements of natural enzymes.(3) Previous experimental studies to explain the structural basis of catalytic powers of enzymes have been largely based on investigation of the binding properties of substrate/substrate analog and transition state/transition-state-analog by enzymes, including examination of related structural changes and interactions. However, there are significant changes in electronic structures and properties of enzyme complexes when substrates (and probably enzymes as well) undergo bond breaking and making processes of enzyme-catalyzed reactions, leading to significant perturbations in their structures, dynamics as well as interactions. Thus, efforts may need to be made to go beyond the investigations of ligand binding properties and to identify new strategies used by enzymes for catalysis based on studying detailed processes of enzyme-catalyzed reactions and cooperative actions of residues in optimizing catalytic efficiency.

One of the well accepted strategies for enzyme catalysis is that enzymes are able to position their residues precisely in enzyme-substrate complexes, especially those near the active sites, and form the reactive conformations for reaction steps they catalyze.(3, 4) The formation of the reactive conformations can also serve as the basis for transition state stabilization.(5) The residues of enzymes may be pre-organized into catalytic competent conformations during the folding process, and a complimentary structural fit between substrate and enzyme (e.g., through the lock and key type of binding)(6, 7) may be involved in the formation of the reactive enzyme-substrate complex. Alternatively, the reactive conformations may be generated through induced fit processes.(7, 8) In the induced fit process, the binding of a substrate may induce a change in the three-dimensional relationship of the active site residues of enzyme, and precise orientation of catalytic groups required for enzyme’s action can be generated through such process (Figure 1a). For the induced fit process, there may be some active-site residues that occupy similar positions before and after substrate binding (e.g., anchor residues), in addition to the residues that undergo large induced-fit conformational changes upon substrate binding (e.g., latch residues) in the formation of the reactive conformations. Previous studies have shown that for protein-protein interactions a lock-and-key mechanism from anchor residues may be followed by a induced-fit process involving latch residues,(9) and such step by step processes may also exist for protein-substrate interactions(10) and for the formation of the reactive conformations for enzyme-substrate complexes. In additional to the conformational changes of enzymes induced by substrates, enzymes may also induce structural distortions or conformational changes of substrates, with the substrate structures moved along the reaction coordinate toward the transition state for the enzyme-catalyzed reactions.(11–13) Furthermore, previous studies have indicated that the reactive conformations of enzyme-substrate complexes may also be generated through the processes of conformational selection and its extension.(10, 14, 15) The importance of formation of the reactive conformations of enzyme-substrate complexes for catalysis has been well recognized,(3, 5) and considerable efforts have been made to design proteins with well-positioned residues through, for instance, computational and machine learning approaches.(16, 17) In addition to the properties of the enzyme-substrate complexes, understanding the interactions between enzymes and transition state is also of crucial, as transition state stabilization is believed to be one of the key factors for enzyme catalysis. Transition state analogs have been extensively used to probe the related properties for transition state stabilization.(1, 18) Nevertheless, transition state analogs are not perfect(19) and may not represent the true transition states that are in the process of bond breaking and making during enzyme-catalyzed reactions.

**Figure 1.**
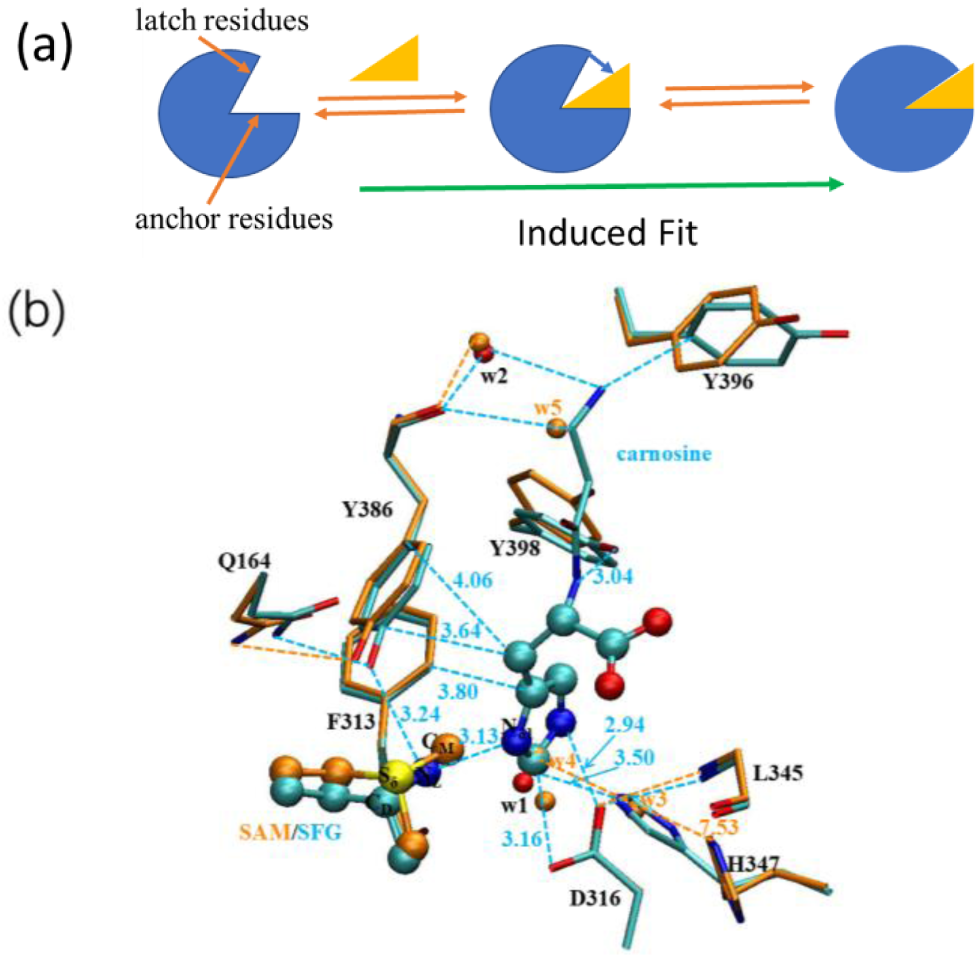
(a) Induced fit model (IF). Blue: enzyme; Yellow: substrate. (b) Superposition of apo enzyme (5YF0, orange) and enzyme-substrate complex (5YF1 with SAM replaced by SFG, light blue). The χ(C-C_α_-C_β_-C_γ_) dihedral angle changes from 55° in 5YF0 to 157° in 5YF1, and r(Carnosine-Cε1···H347-Cε1) decreases from about 7.5Å to 3.5Å accordingly. Part of the SFG/SAM, carnosine and water molecules are shown in ball and stick, and the rest of the active sites are shown in stick. Water molecules from 5YF1 are colored red, and those from 5YF0 in orange. Some distances are also given (in angstroms).

Although the formation of reactive conformations for enzyme-substrate complexes is a crucial and early step for enzyme catalysis, generation of catalytic competent orientations/conformations of residues relative to substrates alone may not be sufficient to lead to efficient catalysis. In many cases, the catalytic competent orientations/conformations of enzyme groups relative to substrate may need to be maintained (probably with strengthened interactions) at least for a part of the reaction process (e.g., going beyond transition state) so that catalytic residues can exert their effects, including transition state stabilization. The formation of the reactive conformations for the substrate complexes does not guarantee that such reactive conformations can be maintained during the enzyme-catalyzed reactions due, for instance, to significant changes in electronic structures and dynamic properties during the bond breaking and making processes. Pre-mature collapse of the reactive conformations can lead to the loss of enzyme’s ability to stabilize transition state, for instance, and significant reduction of catalytic efficiency. To the best of our knowledge, the ability of enzymes to maintain reactive conformations during the reactions has not been considered as an important factor for high catalytic efficiency of enzymes. In this study, QM/MM MD and potential of mean force (PMF) simulations were performed for CARNMT1 to determine the importance of preserving the reactive conformations during the enzyme-catalyzed reaction for efficient catalysis. QM/MM approaches used in this study are especially suitable for studying reaction processes to obtain structural, dynamic and energetic information on enzyme-catalyzed reactions.

CARNMT1 studied in this work is a histidine N_δ1_ position-specific methyltransferase and catalyzes the conversion of carnosine to anserine. The crystal structures for the apo enzyme and enzyme-substrate complex have been determined (PDB IDs: 5YF0 and 5YF1, respectively).(20) CARNMT1 is an ideal system for studying the ability of enzymes to preserve the catalytic competent reactive conformations during catalysis. This is because His347 of the enzyme undergoes large induced-fit conformational changes upon substrate binding and product formation and is the key residue for generation of the reactive conformation of the enzyme-substrate complex for CARNMT1. This property of CARNMT1 makes it easier to study the stability of the reactive conformation during the enzyme catalyzed reaction by monitoring the conformation of His347 with some other active-site residues (see below) being mutated. Moreover, the conformational change of His347 mainly involves the rotation around its C_α_-C_β_ bond with the corresponding χ(C-C_α_-C_β_-C_γ_) dihedral angle changing from about 55° in the apo enzyme to 157° in the enzyme-substrate complex (Figure 1b). The relative activities of several mutants of CARNMT1 compared to that of wild type have been determined along with the structure of the enzyme-product complex (PDB ID: 5YF2).(20) The structures for the apo enzyme and enzyme-product complex are highly similar, and the positions of His347 in 5YF0 and 5YF2 can be superposed well with similar χ(C-C_α_-C_β_-C_γ_) values (55° in 5YF0 and 51° in 5YF2). Thus, the substrate can lead to the induced-fit conformational change involving His347 and generate the reactive enzyme-substrate conformation, while the product (a non-substrate) cannot generate or maintain such conformation.(20)

One of the widely used approaches to understand the origin of enzyme’s catalytic powers and provide important insights for enzyme design is to examine the effects of mutations on enzyme-catalyzed reactions and catalytic efficiencies.(5, 19, 21) We have adopted this approach in this study and examined the ability of the wild-type CARNMT1 and different mutants (including Phe313Ala, Asp316Ala, Tyr386Ala, Tyr396Asp, Tyr398Ala as well as His347Phe) to preserve the reactive conformation during the enzyme-catalyzed reaction; the correlation between this ability and the catalytic efficiency was also studied. The results from our QM/MM MD and free energy (PMF) simulations showed that the wild-type enzyme can preserve the catalytic competent reactive conformation even after the reaction passed the transition state (TS). Thus, the reason that the wild-type enzyme has high catalytic efficiency is not only due to the formation of catalytic competent reactive conformation for the enzyme-substrate complex but also related to the ability of wild type to preserve the catalytic competent conformation during the reaction. The suggestion for the importance of preserving the reactive conformation has been supported by the results on the mutants. Indeed, for the Tyr386Ala, Tyr396Asp and Tyr398Ala mutants the wild-type like reactive conformation (for which the methyl donor and acceptor are arranged similarly as in wild type, and His347 is also in the active “hold” position) can be formed in the mutant-substrate complex. However, the wild-type like reactive conformation for these mutants collapses in the early stage of reaction before the reaction can reach TS. Such pre-mature collapse of the reactive conformation with additional solvent exposure of the active site can lead to diminished TS stabilization and is likely to be responsible for the significant reduction of catalytic efficiency obtained computationally from this work and observed experimentally previously.(20) For the Asp316Ala and Phe313Ala mutants, the wild-type like reactive conformation cannot even be formed in the substrate complex due to the loss of the key interactions that fix the position of the methyl acceptor, imidazole ring on the substrate. The results suggest that generating precise catalytic competent orientations for the reactants and catalytic groups in enzyme-substrate complexes through the well-known models may be a necessary but not sufficient condition for efficient catalysis. Moreover, they showed that one of the reasons for some enzymes to have high catalytic efficiency may be related to their ability to preserve the reactive conformations with well positioned residues during the crucial parts of enzyme-catalyzed reactions. The results on the mutants may have some implications for uncatalyzed reactions as well. Low reactivity for uncatalyzed reactions could also be related to the inability for the reactants to maintain reactive conformations and favorable interactions within the environments during the reaction processes, due probably in part to dynamic features of non-enzymatic environments.

## Results

### The wild-type enzyme has the ability to preserve the reactive conformation beyond TS for efficient catalysis

The average active-site structure of the CARNMT1-SAM-carnosine complex obtained from QM/MM MD simulations is shown in Figure 2a. The interactions between carnosine and the key residues in the active site region (e.g., Asp316, His347, Tyr386, Tyr396, etc.) may play a crucial role in the substrate binding as well as catalysis. The average structure and interactions obtained from the simulations are quite similar to those observed experimentally (Figure 1b).(20) For instance, the two oxygen atoms of the Asp316 side chain interact with the C_ε1_ and N_ε2_ atoms of the substrate imidazole ring, respectively, in both crystal structure and reactant state from simulations; the corresponding average distance to C_ε1_/N_ε2_ is 3.43 Å/2.78 Å obtained from the simulations (Figure 2a), while it is 3.16 Å/2.94 Å in the crystal structure (Figure 1b). Other interactions that are also similar between the average structure obtained from the simulations and crystal structure include the key T-shape “π-π” stacking interactions between the Phe313 ring and carnosine imidazole ring (e.g., 3.95 Å and 3.80 Å as shown in Figure 2a and 1b, respectively) and between the His347 and carnosine imidazole rings (e.g., 3.67 Å and 3.50 Å as shown in Figure 2a and 1b, respectively), the hydrophobic contacts involving the His C_β_ atom of the substrate and the ring of Tyr386 (e.g., 3.73 Å and 3.64 Å in Figure 2a and 1b, respectively), and the hydrogen bond between the hydroxyl group of Tyr398 and the substrate (3.13 Å and 3.04 Å in Figure 2a and 1b, respectively). The positions of the active site residues are fixed by the interactions with the substrate carnosine and their neighboring residues. For instance, the interaction between the backbone carbonyl oxygen of Leu345 and N_δ1_−H of His347 stabilizes the His347 position, and His347 then forms the “π-π” interaction with the imidazole ring of the substrate as mentioned above. We have also observed some interactions involving the transferable methyl group in our simulated structure as a result of replacing SFG with SAM. For example, the hydroxyl group of Tyr386 and the backbone carbonyl oxygen of Phe313 form carbon hydrogen bonds(22) with the transferable methyl group of SAM. In general, the average structure obtained from the QM/MM MD simulations and crystal structure are highly similar; the existence of some small differences is expected as SFG in the crystal structure was replaced by SAM for the simulations. The *r-θ* and *r-β* distribution maps of the enzyme-substrate complex of wild type are also given in Figure 2a (see Methods for the definitions). As can be seen from the distribution maps, there are many structures containing relatively short distances for *r*(C_M_−N_δ1_) (average value of 3.09 Å) with the average *θ* and *β* angles of 21.92° and 157.07°, respectively. These structures formed during the simulations are suitable for the methylation reaction based on the criteria obtained from our earlier studies that methylation reactions can occur efficiently if *r*(C_M_−N_δ1_), *θ* and *β* are close 3 Å, 0°, 180°, respectively.(23, 24) Therefore, when carnosine binds to the active site of the enzyme, it can form the near-attack reactive conformation(25) to facilitate methyl transfer.

**Figure 2.**
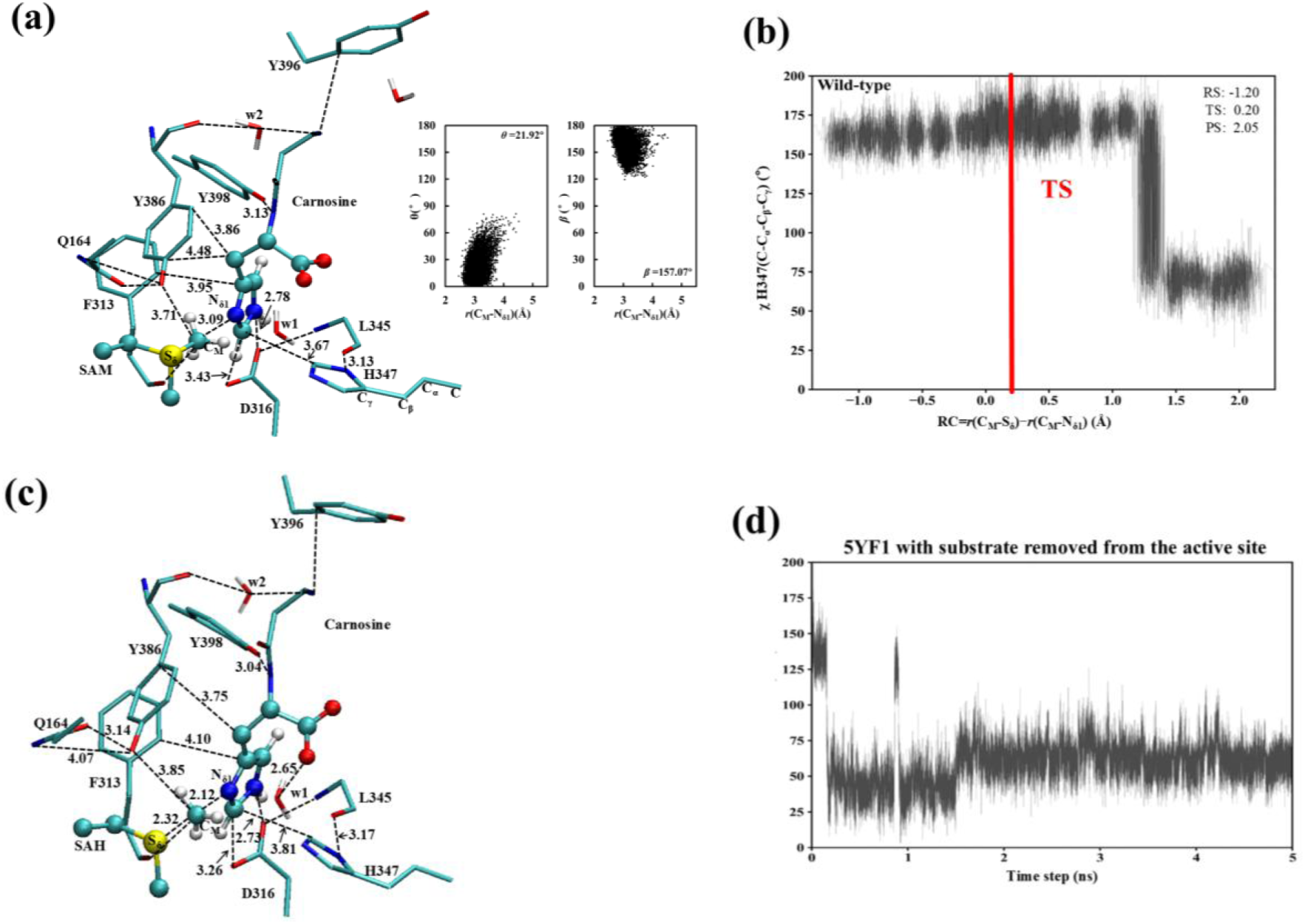
(a) The average structure for the active site of the enzyme-substrate complex containing SAM and carnosine obtained from the QM/MM MD simulations in wild-type CARNMT1 (left) along with the *r*(C_M_-N_δ1_)/*θ* and *r*(C_M_-N_δ1_)/*β* distribution maps (right). The average values of *r*(C_M_-N_δ1_), *θ* and *β* are 3.09Å, 21.92° and 157.07°, respectively. The χ(C-C_α_-C_β_-C_γ_) dihedral angle of His347 is about 160°, and *r*(carnosine-C_ε1_···H347-C_ε1_) is 3.7Å. (b) The fluctuations of the χ(C-C_α_-C_β_-C_γ_) dihedral angle of His347 in the PMF windows along the reaction coordinate. The red vertical bar designates the location of near transition state for the methyl transfer. (c) The average structure for the active site near transition state obtained from the QM/MM PMF simulations for methylation in wild-type CARNMT1. (d) The χ(C-C_α_-C_β_-C_γ_) values from the simulations after the substrate was first removed from the active site in the enzyme-substrate complex.

Figure 2b shows the fluctuations of the χ(C-C_α_-C_β_-C_γ_) dihedral angle of His347 collected from each of the PMF windows along the reaction coordinate from the QM/MM free energy (PMF) simulations for the methyl transfer process (see Methods). As is shown in Figure 2b, the wild-type enzyme can preserve the catalytic competent reactive conformation with His347 in the active “hold” position [i.e., χ(C-C_α_-C_β_-C_γ_) ∼ 160° as observed in the crystal structure](20) well after the reaction passes TS [R = *r*(C_M_−S_δ_) − *r*(C_M_−N_δ1_) ∼ 0.2Å]. The reactive conformation collapses at R = *r*(C_M_−S_δ_) − *r*(C_M_−N_δ1_) ∼ 1.2 Å with χ(C-C_α_-C_β_-C_γ_) of His347 changing from the active “hold” position to the inactive “released” position (i.e., χ(C-C_α_-C_β_-C_γ_) ∼ 60°), consistent with the crystal structure of the product complex.(20) The suggestion for the ability of wild-type to maintain the reactive conformation during the key part of the reaction is supported by the average structure of the active site near transition state of the methylation reaction obtained from the QM/MM PMF simulations (Figure 2c). Indeed, Figure 2c shows that the reactive conformation of the CARNMT1-substrate complex (Figure 2a) is well preserved near the transition state with only some small modifications. For instance, the T-shape “π-π” stacking interactions between the imidazole ring of the substrate and the Phe313/His347 side chain remain intact near the transition state that help to generate the well-positioned interactions and protect the substrate from the exposure to solvent. We have also examined what would happen if the substrate in the reactive conformation of the enzyme-substrate complex was removed from the active site. Figure 2d shows that if the substrate was removed, His347 would move quickly from the active “hold” to the inactive “released” position (as observed in the apo enzyme. See Figure 1b).(20) The ability to predict the conformational changes of His347 for the product formation and in the apo enzyme indicates that the QM/MM MD simulations are reasonably reliable.

### Tyr386Ala, Tyr396Asp and Tyr398Ala mutations lead to the loss of the enzyme’s ability to preserve the reactive conformation in early stage of the reaction

The average active-site structures of the Tyr386Ala-substrate, Tyr396Asp-substrate and Tyr398Ala-substrate complexes obtained from the QM/MM MD simulations are given in Figure 3a, 3c and 3e, respectively, along with the corresponding *r-θ* and *r-β* distribution maps. For Tyr386Ala, some interactions involving Tyr386 observed in wild type are lost in the mutant complex, as expected. For instance, the carbon hydrogen bond between the hydroxyl oxygen of Tyr386 and transferable methyl group and the hydrophobic contacts involving the His C_β_ atom of the substrate and the ring of Tyr386 are lost. Almost all the other interactions remain similar as those observed in wild type with only small derivations. Figure 3a show that the average structure of the Tyr386Ala-substrate complex is in the wild-type like reactive conformation with His347 in the active “hold” position (χ(C-C_α_-C_β_-C_γ_) ∼ 160°). Moreover, there is a large population of structures from the MD simulations with relatively short *r*(C_M_−N_δ1_) distances (average value of 3.28Å) and good *θ* and *β* values for methylation (average values of 27.15° and 162.54°, respectively), i.e., the near attack conformations are highly populated. The fluctuation of the χ(C-C_α_-C_β_-C_γ_) dihedral angle of His347 along the reaction coordinate obtained from the PMF simulations is plotted in Figure 3b for Tyr386Ala. As is evident from Figure 3b, the χ(C-C_α_-C_β_-C_γ_) value changed from about 160° to 60° in early stage of the methyl transfer reaction [R = *r*(C_M_−S_δ_) − *r*(C_M_−N_δ1_) ∼ –1.5 Å] well before the reaction could reach the transition state [R = *r*(C_M_−S_δ_) − *r*(C_M_−N_δ1_) ∼ 0.2Å]. Thus, the wild-type like reactive conformation of the catalytic groups relative to the substrate in the Tyr386Ala-substrate complex could not be preserved during the reaction to allow catalytic residues to fully exert their catalytic effects. The pre-mature collapse of such conformation is expected to affect the efficiency of methylation process, contributing to an increase of free energy barrier (see below) and decrease of the activity.(20) Figure 3c and 3e show that for the Tyr396Asp and Tyr398Ala mutants, the wild-type like reactive conformations with His347 in the active “hold” position also exist in the mutant-substrate complexes; there are also large populations of structures with relatively short *r*(C_M_−N_δ1_) distances (average values of 3.07 Å and 3.14 Å for Tyr396Asp and Tyr398Ala, respectively) as well as good *θ* values (average values of 26.1° and 19.72° for Tyr396Asp and Tyr398Ala, respectively) and good *β* values (average values of 163.67° and 168.34° for Tyr396Asp and Tyr398Ala, respectively) for methylation. Thus, the near attack conformations are highly populated in these two cases as well. The interactions with the substrate are also quite similar to those observed in wild type with the exception of those involved the mutated residues (i.e., Asp396 and Ala398 for Tyr396Asp and Tyr398Ala, respectively). Nevertheless, as is evident from Figures 3d and 3f, the both wild-type like reactive conformations observed in the mutant-substrate complexes collapsed at R = *r*(C_M_−S_δ_) − *r*(C_M_−N_δ1_) ∼ –0.8 Å and –1.0 Å, respectively. Thus, like the case of Tyr386Ala, the wild-type like reactive conformations for these two mutants could not be preserved, contributing to an increase of free energy barrier and decrease of the activity.

**Figure 3.**
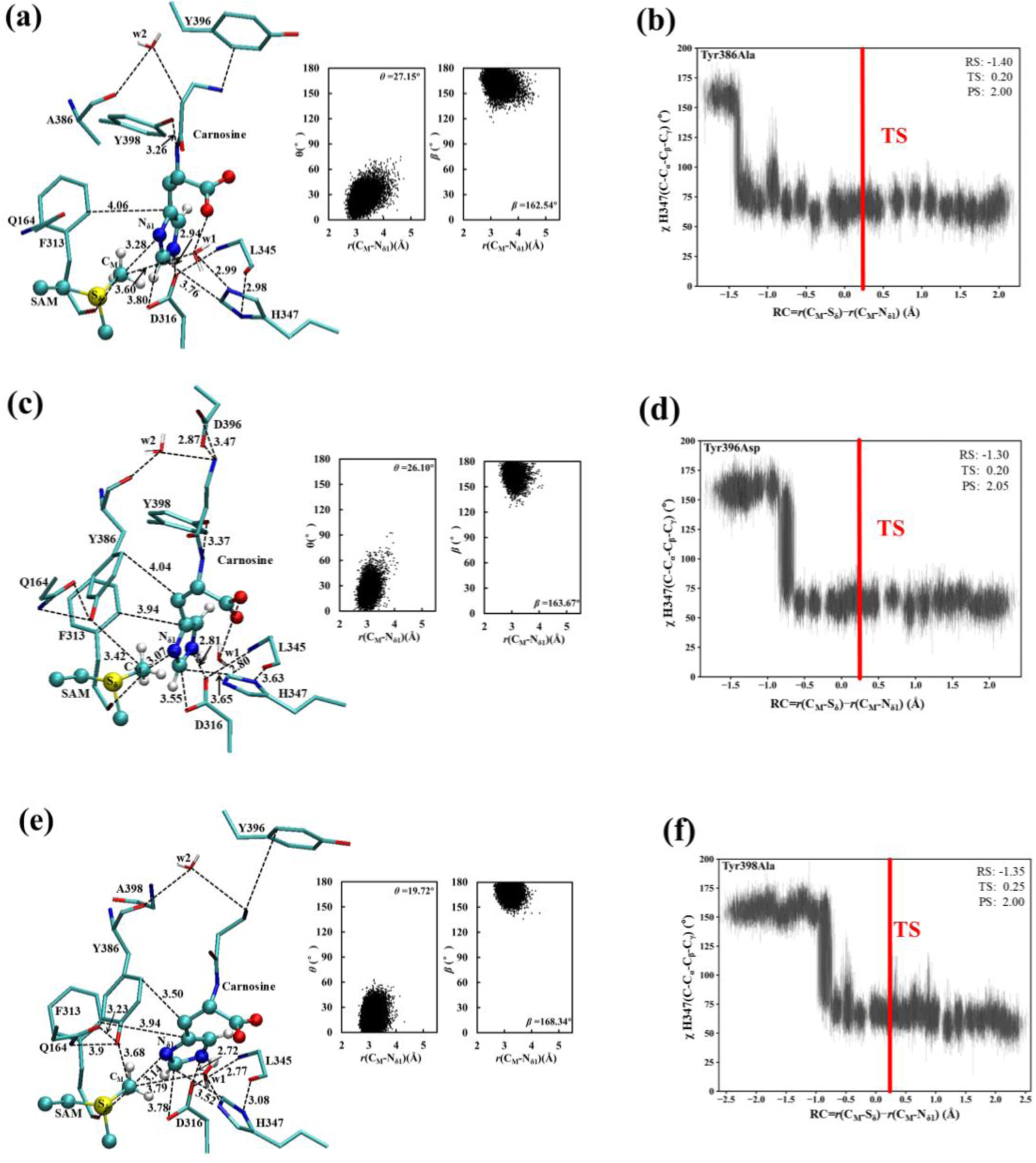
(a), (c) and (e) The average structures for the active sites of the Tyr386Ala-substrate, Tyr396Asp-substrate and Tyr398Ala-substrate complexes, respectively, along with the corresponding distribution maps. (b), (d) and (f) The fluctuations of the χ(C-C_α_-C_β_-C_γ_) dihedral angle of His347 in the PMF windows along the reaction coordinate for the Tyr386Ala, Tyr396Asp and Tyr398Ala complexes, respectively.

The average structures for the active sites near transition state of methylation for the Tyr386Ala, Tyr396Asp and Tyr398Ala complexes are given in Figure 4a, 4b and 4c, respectively. Consistent with the results in Figure 3b, 3d and 3f, the wild-type TS conformation with His347 in the active “hold” position (Figure 2c) does not exist near transition state for these three mutants; His347 is now in the inactive “released” position. Figure 4a, 4b and 4c also show that after His347 was changed to the inactive “released” position during the reaction, additional water molecule moved into the active site and occupied the original position of His347 in the mutants-substrate complexes (see Figures 3a, 3c and 3e). The same water molecule also exists in the apo crystal structure (w3 in 5YF0. See Figure 1b) and in the product complex (5YF2). Previous studies on other enzymes (e.g., ketosteroid isomerase) have indicated that water molecules are less effective in stabilizing transition state compared to the well positioned active-site residues.(26, 27) Thus, the change of the His347 position and exposure of the transition state to additional solvent molecule(s) for the mutants is expected to lead to diminished TS stabilization and be responsible for the reduction of catalytic efficiency obtained computationally from this work (see below) and observed experimentally previously.(20)

**Figure 4.**
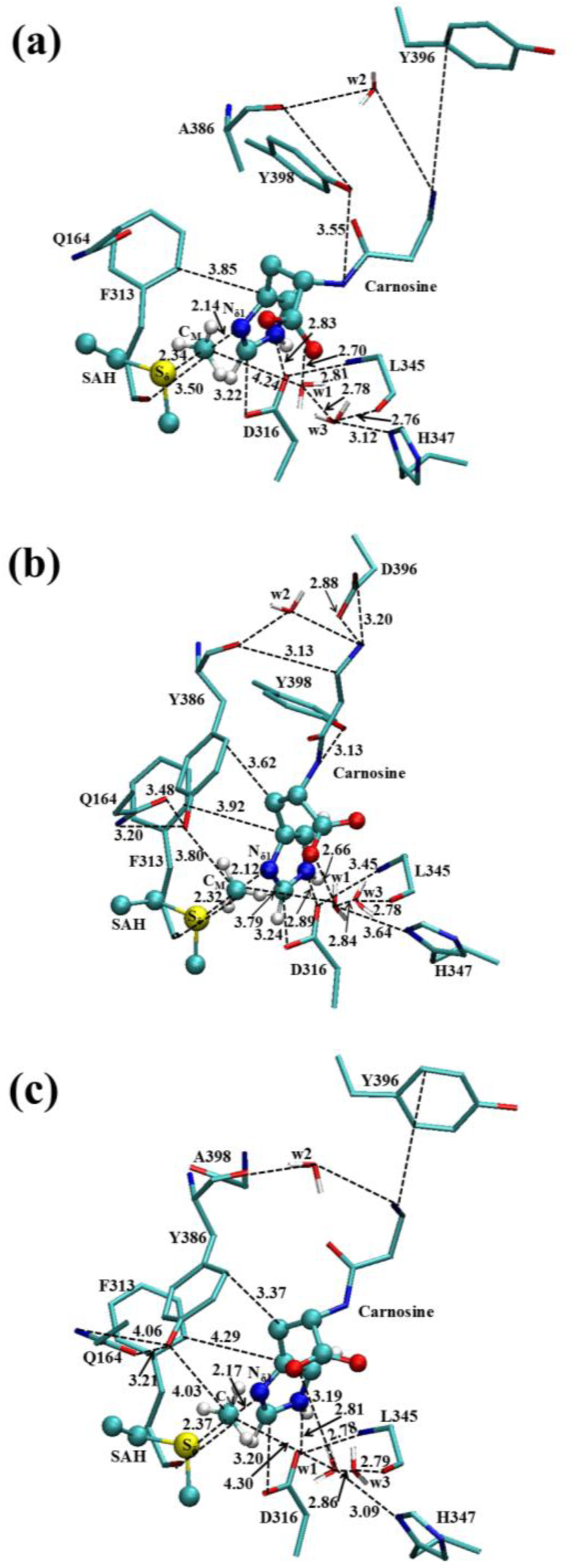
(a), (b) and (c) The average structures for the active sites near transition state of methylation for the Tyr386Ala, Tyr396Asp and Tyr398Ala complexes, respectively.

It would be of interest to examine the effect of replacing His347 by Phe on the preservation of the reactive conformation. The average structure and distribution maps for the His347Phe-substrate complex are given in the Supplementary Materials (Figure S1). As in the cases of the Tyr386Ala, Tyr396Asp and Tyr398Ala mutants, the His347Phe-substrate complex also has a large population of structures with the relatively short r(C_M_−N_δ1_) distances and good θ and *β* values for methylation. Nevertheless, the reactive conformation also collapsed before the reaction reached the TS [at R = r(C_M_−S_δ_) − r(C_M_−N_δ1_) ∼ –0.2 Å]; this is consistent with our PMF results (see below) and the experimental observation(20) that more than 80% of the activity was lost due to the His347Phe mutation.

### Inability to form the reactive conformation in the mutant-substrate complex for Asp316Ala and Phe313Ala

The results given in Figure 2a and 2c and previous experimental data(20) support the suggestion that Asp316 stabilizes the conformation of the N_ε2_-H tautomeric form of the substrate and makes it possible for the transferable methyl group of SAM to point towards the electron lone pair of N_δ1_ of carnosine prior to the reaction. The Asp316 residue in the CARNMT1 seems to play a similar and presumably more important role as the Asn255 residue in SETD3(23). If the hydrogen bonding/carbon hydrogen bonding interactions between Asp316 and carnosine are removed (e.g., through mutations), the binding of the correct tautomer and the formation of the near-attack reactive state are likely to be compromised, leading to a significant reduction or loss of the activity. The average active-site structure of the Asp316Ala-substrate complex obtained from the simulations is given in Figure 5a along with the distribution maps. Although other active site residues (e.g., Phe313, His347, Leu345 and Tyr386) may also be important for anchoring the N_ε2_-H tautomeric form and the formation of the reactive conformation, they cannot achieve this by themselves without Asp316. Indeed, the distribution maps in Figure 5a show that the population of the structures with relatively short *r*(C_M_−N_δ1_) distances along with good *θ* and β values for methylation become much smaller in the Asp316Ala mutant compared to the case of wild-type (Figure 2a); e.g., the average values of *r*(C_M_−N_δ1_), *θ* and *β* have changed from 3.09 Å, 21.92° and 157.07° in Figure 2a to 3.45 Å, 49.34° and 151.89° in Figure 5a, respectively. Moreover, comparison of the structures of Figure 2a and Figure 5a shows that the His residue of the substrate underwent a rotation in Asp316Ala, and this conformational change of the substrate also affects the correct alignment between the methyl donor and acceptor. Furthermore, the imidazole ring of His347 flipped away from the active “hold” position in Figure 2a and now has a χ(C-C_α_-C_β_-C_γ_) value of about 60° (Figure 5a and 5b). Thus, His347 is already in the inactive “released” position in the Asp316Ala-substrate complex even before the reaction starts. The fluctuations of the χ(C-C_α_-C_β_-C_γ_) dihedral angle of His347 along the reaction coordinate obtained from the PMF simulations are given in Figure 5b. As is evident from Figure 5b, the χ(C-C_α_-C_β_-C_γ_) values remained around 60° during the methyl transfer reaction, and the reactive conformation could not be generated during the reaction. The loss of such reactive conformation in the substrate complex as well as during the reaction is expected to affect the efficiency of methylation process significantly. The average structure for the active site of the Phe313Ala-substrate complex is given in Figure 5c. As is evident from Figure 5c, His347 is already in the inactive “released” position in the Phe313Ala-substrate complex, and similar to the case of Asp316Ala, the reactive conformation could not be generated during the methylation reaction.

**Figure 5.**
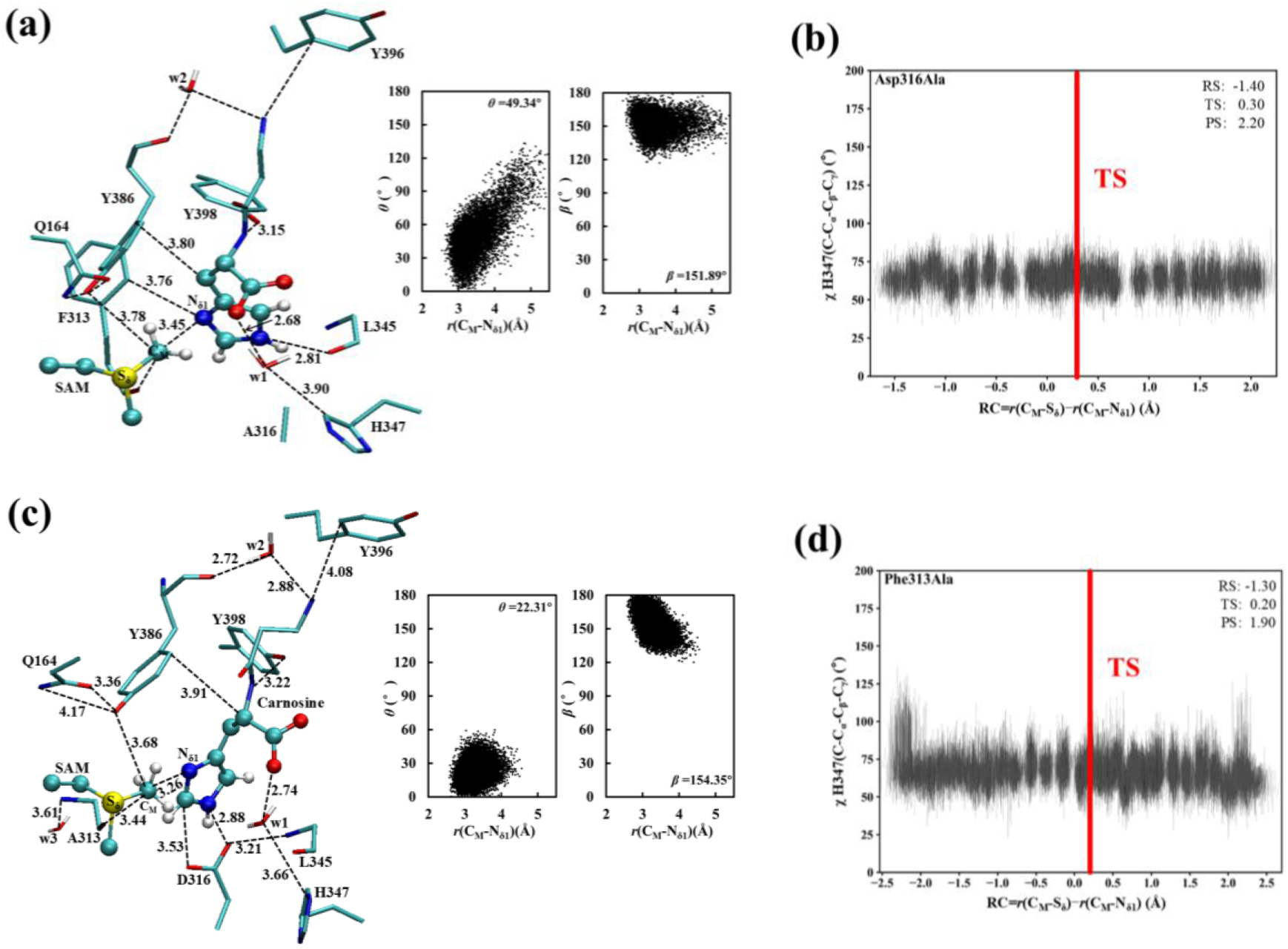
(a) and (c) The average structures for the active sites of the Asp316Ala-substrate, and Phe313Ala-substrate complexes, respectively, along with the corresponding distribution maps. (b) and (d) The fluctuations of the χ(C-C_α_-C_β_-C_γ_) dihedral angle of His347 in the PMF windows along the reaction coordinate for the Asp316Ala and Phe313Ala complexes, respectively.

### The free energy profiles for methylation in wild-type and some mutants

The free-energy (PMF) profiles for the methylation reaction in the wild-type CARNMT1, Phe313Ala, Asp316Ala, Tyr386Ala, Tyr396Asp, Tyr398Ala and His347Phe as a function of the reaction are given in Figure 6. As is shown in Figure 6, the free energy barriers increase by 2.5, 3.72, 3.42, 1.71, 1.78 and 1.14 kcal·mol^-1^ relative to that of wild type as a result of the Phe313→Ala, Asp316→Ala, Tyr386→Ala, Tyr396→Asp, Tyr398→Ala and His347→Phe mutations, respectively. The increases of the free energy barriers are consistent with the decreases of the activity observed in the previous experimental observations.(20) Interestingly, the increases of the free-energy barriers compared to that of wild-type seem to have some correlations with the ability/inability of the mutants to preserve the wild-type like reactive conformation for most of the cases. For instance, the Tyr396Asp mutant had its wild-type like reactive conformation collapsed [at R = ∼ –0.8 Å] later than the Tyr386Ala mutant [at R = ∼ –1.5 Å] (i.e., the ability of Tyr396Asp to preserve the wild-type like reactive conformation is better than that of Tyr386Ala), while its free energy barrier (18.3 kcal/mol) for the reaction is lower than that of Tyr386Ala (∼ 20 kcal/mol). For Tyr396Asp and Tyr398Ala, the wild-type like reactive conformations collapsed at the similar positions along the reaction coordinate (i.e., at R = *r*(C_M_−S_δ_) − *r*(C_M_−N_δ1_) ∼ –0.8 Å and –1.0 Å for Tyr396Asp and Tyr398Ala, respectively), the free energy barriers are also similar (18.30 and 18.37 kcal/mol, respective). The Asp316Ala mutant has the least ability to preserve the wild-type like reactive conformation among the mutants studied here, and its free energy barrier for methylation is also the highest. Although the experimental k_cat_ values for the mutants are not available for comparison, the general trends obtained are consistent with the relative activities of the mutants measured previously.(20) For instance, Phe313Ala, Asp316Ala and Tyr386Ala were found to be among the least active mutants from the earlier study,(20) while they have the relatively high free energy barriers from this study. The free energy barrier (17.73 kcal/mol) for His347Phe is the lowest among the mutants studied, and it was also found to be the most active among the mutants.(20)

**Figure 6.**
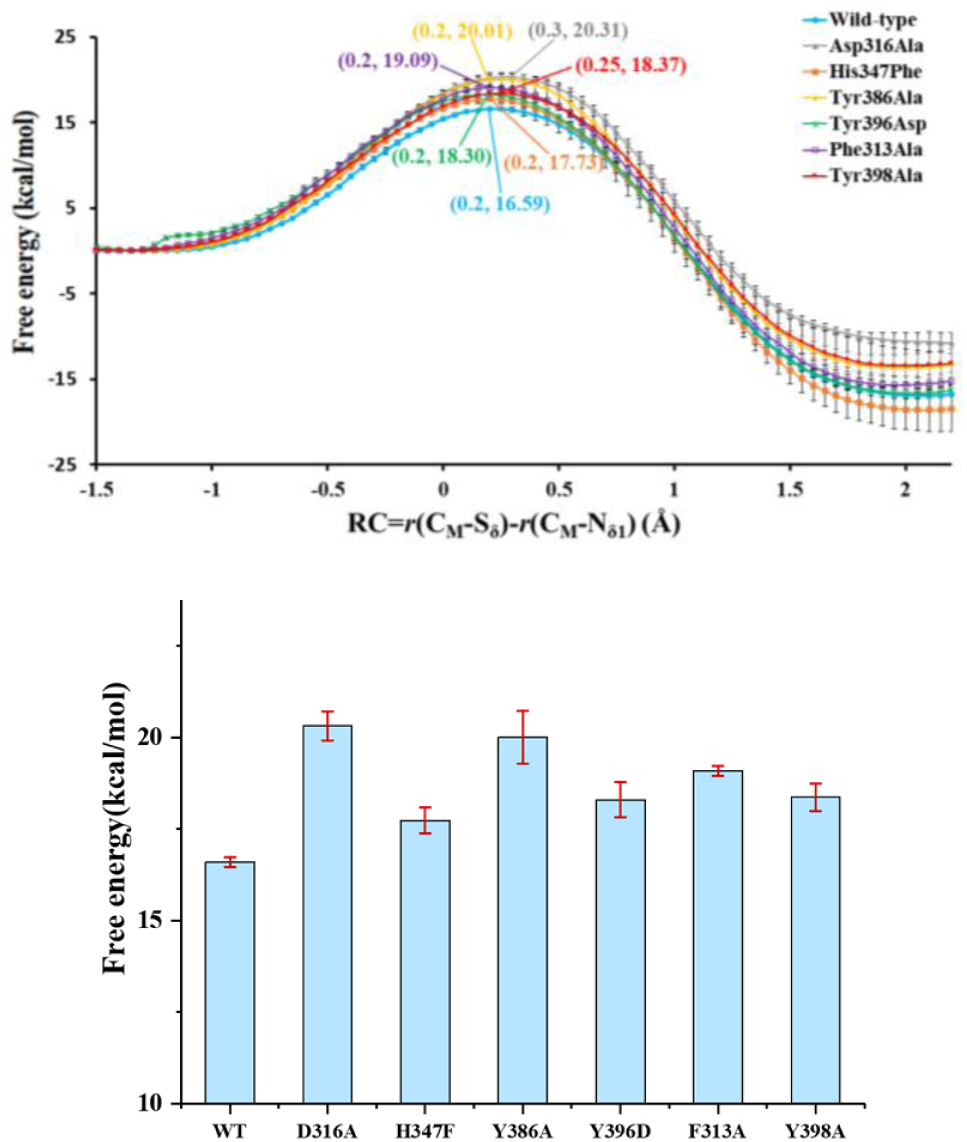
Free energy (PMF) profiles for the methylation reaction in wild-type CARNMT1, Phe313Ala, Asp316Ala, Tyr386Ala, Tyr396Asp, Tyr398Ala and His347Phe, respectively, as a function of the reaction coordinate [R = *r*(C_M_−S_δ_) − *r*(C_M_−N_δ1_)]. Wild type: blue line with the free energy barrier of 16.59 kcal/mol; Phe313Ala: purple line with the free energy barrier of 19.09 kcal/mol; Asp316Ala: grey line with the free energy barrier of 20.31 kcal/mol; Tyr386Ala: yellow line with the free energy barrier of 20.01 kcal/mol; Tyr396Asp: green line with the free energy barrier of 18.30 kcal/mol; Tyr398Ala: red line with the free energy barrier of 18.37 kcal/mol; His347Phe: orange line with the free energy barrier of 17.73 kcal/mol.

## Discussion

Considerable efforts have been made to understand the strategies used by enzymes for catalysis and to provide important insights for enzyme design,(1, 2) but our understanding of the mechanisms by which enzymes achieve enormous rate enhancements is still not complete. One of the wildly accepted strategies used by enzymes is that enzymes can position catalytic residues precisely in enzyme-substrate complexes and form reactive conformations for catalysis (i.e., catalysis by positioning). However, the catalytic competent conformations of enzyme systems also need to be maintained during crucial parts of enzyme-catalyzed reactions to allow catalytic residues to exert the effects. The existence of the reactive conformations for enzyme-substrate complexes does not guarantee that the catalytic competent conformations would also be present during reactions. Nevertheless, little attention has been paid to the importance of preserving the reactive conformations during reactions, presumably due in part to the difficulty in observing the reaction processes experimentally.

In this study, we performed QM/MM MD and free energy simulations on CARNMT1 and its mutants to determine the importance and requirements for preservation of catalytic competent reactive conformations during enzyme-catalyzed reactions. We have compared the ability of the wild-type CARNMT1 and different mutants (including Phe313Ala, Asp316Ala, Tyr386Ala, Tyr396Asp, Tyr398Ala as well as His347Phe) to preserve the reactive conformation during the enzyme-catalyzed reaction through monitoring the conformation of His347. The key conclusions of this work are given in Figure 7. For the wild-type enzyme, the catalytic competent conformation could not only be formed in the enzyme-substrate complex but also be preserved well after the reaction passed TS. This allows the catalytic residues to exert their catalytic effects. For the Type 1 mutants, including Tyr386Ala, Tyr396Asp and Tyr398Ala, the wild-type like reactive conformations could be formed in the mutant-substrate complexes but collapsed in the early stage of reaction, leading to diminished TS stabilization and significant reduction of catalytic efficiency. Thus, generating precise orientations of catalytic groups in enzyme-substrate complexes through the well-known models may be necessary but not sufficient for efficient catalysis. For the Type 2 mutants, including Asp316Ala and Phe313Ala, the reactive conformation could not even be formed in the mutant-substrate complexes with relatively more significant reductions of the activity. One of the key reasons that we have chosen CARNMT1 for this study is because of the simplicity and clarity for monitoring the formation and collapse of the reactive conformation during the methylation reaction for the Type 1 mutants. For some other enzymes, deviations from the potentially best reaction paths and interactions (e.g., as occurred, for instance, for the wild-type CARNMT1) may involve many subtle changes in structures during enzyme-catalyzed reactions. Nevertheless, the principle obtained here may hold for some other enzymes as well, and for these enzymes one of the reasons they have high catalytic efficiency may be related to their ability to preserve the reactive conformations during crucial parts of enzyme-catalyzed reactions.

**Figure 7.**
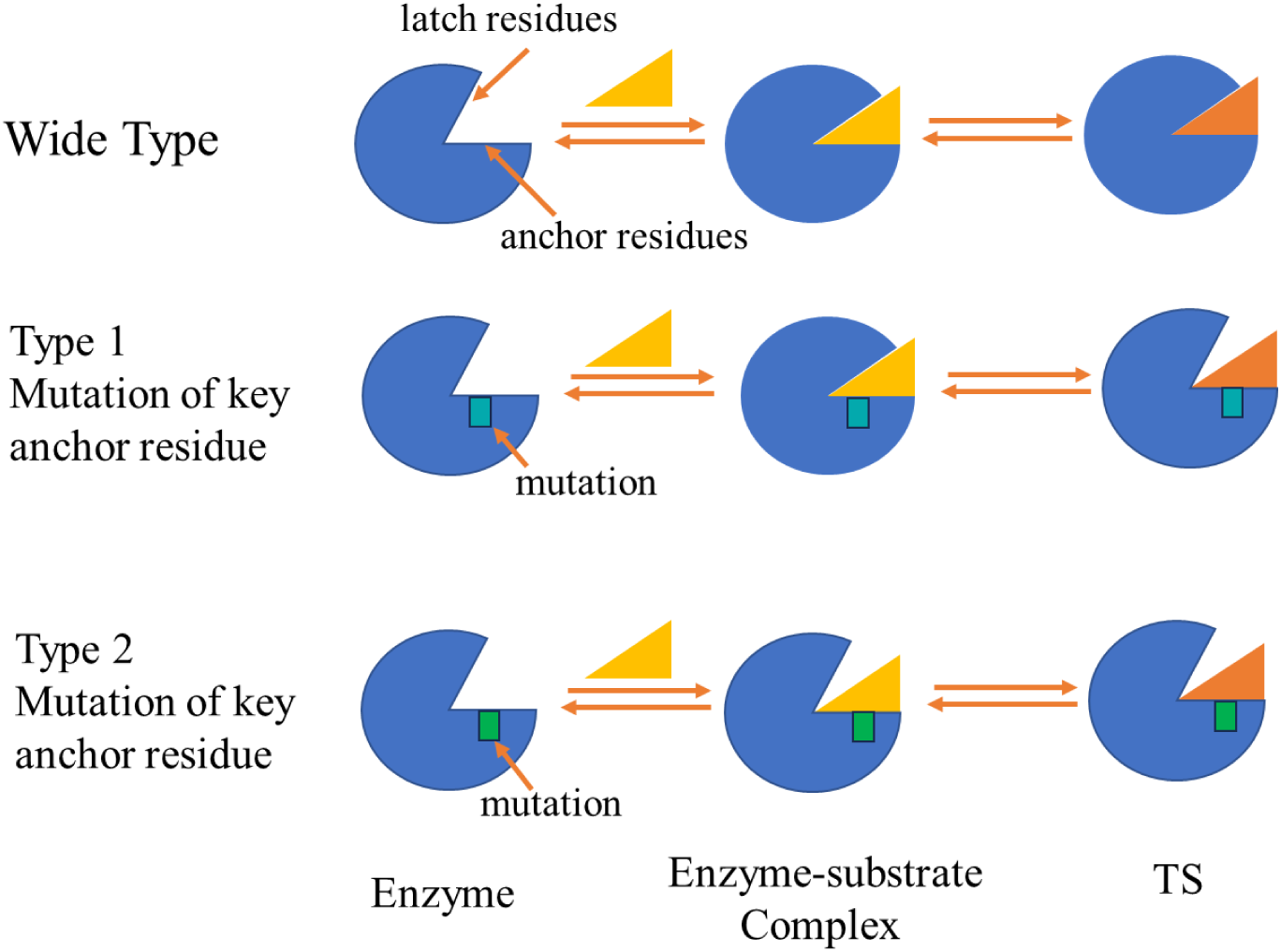
Wild-type enzymes can preserve catalytic competent induced-fit configuration at TS. Type 1 mutation of key anchor residue (e.g., Tyr386Ala) leads to the collapse of the reactive conformation before the reaction reaches TS, even though such conformation can be formed in the enzyme-substrate complex. Type 2 mutation of key anchor residue (e.g., Asp316Ala) destroys the enzyme’s ability to form the reactive conformation with the substrate.

The results on the mutants may have some implications for uncatalyzed reactions. For uncatalyzed reactions, low reactivity could be due to the inability for the reactants to form reactive configurations and/or favorable interactions with the environment before reaction starts (e.g., like the Type 2 mutants). However, low reactivity might also be related to the failure of the reactants to maintain reactive configurations and/or favorable interactions with the environment during reaction processes (e.g., like the Type 1 mutants), due probably to dynamic features of non-enzymatic environments. The principle discussed here may have some implications for the design of artificial catalysts, such as catalytic antibodies and engineered proteins.

## Materials and Methods

The CHARMM program(28, 29) was used to perform the QM/MM MD and free energy simulations to study the formation and collapse of the reactive conformation in the enzyme-substrate complex and during the catalysis. The effects of the Phe313→Ala, Asp316→Ala, Tyr386→Ala, Tyr396→Asp, Tyr398→Ala and His347→Phe mutations on the stability of the wild-type like reactive conformations for the substrate complexes and during the N_δ1_ methylation were also determined. The crystal structure of CARNMT1 (PDB ID: 5YF1) containing carnosine and SFG(20) and was used to create the reaction state structure of the enzyme-substrate complex, and the N and C atoms of SFG was manually changed to C and S atoms, respectively, to generate SAM. The crystal structure of the apo enzyme (PDB ID: 5YF0) containing SAM was also used for the simulations (Figure S2). The simulations were also performed for the case where carnosine was removed from 5YF1 in the initial structure (with SFG changed to SAM). This allows testing whether His347 could move back to its position in the apo enzyme when the substrate was removed from the enzyme-substrate complex. To perform the QM/MM MD simulations on the mutants, the Phe313→Ala, Asp316→Ala, Tyr386→Ala, Tyr396→Asp, Tyr398→Ala or His347→Phe mutation was performed manually in the crystal structure of the wild type (5YF1).

The −CH_2_−CH_2_−S^+^(Me)−CH_2_− part of SAM and the whole substrate were treated by QM, and the rest of the system was treated by MM. The QM region and MM region were separated by using the link-atom approach(30) implemented in CHARMM; the QM region contains a total of 44 atoms, with the net charge of +1. The stochastic boundary molecular dynamics method(31) was used for the QM/MM MD and PMF simulations. A modified TIP3 water model(32) was used for the solvent. The DFTB3 method(33–36) was applied to the QM atoms, and the CHARMM potential function (PARAM27)(37) was adopted for the MM atoms. Hydrogen atoms of the system were built with the HBUILD module(38) in CHARMM. The reference center assigned by the system was chosen to be the N_δ1_ atom of target substrate carnosine. The system was separated into a reaction zone and a reservoir region, and the reaction zone was further divided into a reaction region (a sphere with radius r of 20 Å) and a buffer region (20 Å ≤ r ≤ 22 Å). The whole system contained around 5700 atoms, including about 460 water molecules. The initial structures were optimized for the entire stochastic boundary systems with the steepest descent (SD) and adopted-basis Newton−Raphson (ABNR) methods. The systems were gradually heated from 50.0 to 310.15 K in 50 ps. The time step for the integration of equation of motion was 1 fs, and the coordinates were saved every 50 fs for analyses. For each of the systems, the QM/MM MD simulations of 3 ns were carried out. The distribution maps of *r*(C_M_-N_δ1_)/*θ* and *r*(C_M_-N_δ1_)/*β* (see Figure 8 for definitions) were generated in each case and the fluctuations of the χ(C-C_α_-C_β_-C_γ_) dihedral angle and r(carnosine-C_ε1_···H347-C_ε1_) distance of His347 were monitored.

**Figure 8.**
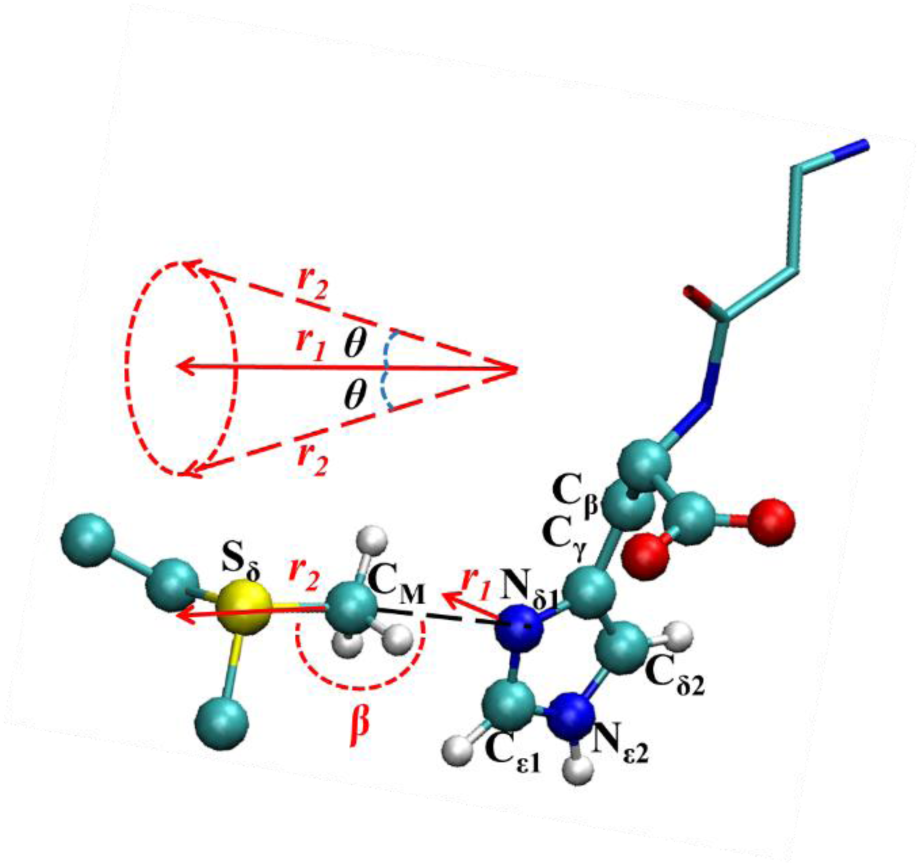
Relative orientation of SAM and carnosine in the reactant complex. Here, *θ* is defined as the angle between two vectors r_1_ and r_2_, which represent the direction of the C_M_−S_δ_ bond and the direction of the lone pair of electrons on N_δ1_, respectively. r(C_M_−N_δ1_), *θ* and β(S_δ_−C_M_−N_δ1_) angle together can provide important information for the formation of the reactive conformation for the enzyme (mutant)-substrate complex.

The umbrella sampling method(39) was applied along with the weighted histogram analysis method (WHAM)(40) in the determination of the free energy change along the reaction coordinate for the methyl transfer from S-adenosyl-L-methionine (SAM) to the N_δ1_ atom of carnosine. The reaction coordinate was defined as a linear combination of *r*(C_M_−N_δ1_) and *r*(C_M_−S_δ_) [R = *r*(C_M_−S_δ_) − *r*(C_M_−N_δ1_)]. For the methyl transfer process, 28 windows were used, and for each window, the production runs of 100 ps were performed after 50 ps equilibration. The force constants of the harmonic biasing potentials used in the PMF simulations were 50 to 400 kcal·mol^−1^·Å^−2^. In each case, four independent PMF simulations were performed. The free energies and statistical errors were taken as the average values and standard deviations from the four runs, respectively. The fluctuations of the χ(C-C_α_-C_β_-C_γ_) dihedral angle as well as r(carnosine-C_ε1_···H347-C_ε1_) distance of His347 were monitored in each PMF window along the reaction coordinate to detect when the reaction conformation was formed or collapsed during the methyl transfer reaction.

## Author Contributions

P.Q. and H.G. designed research; M.Y., H.D. A.R., P.Q. and H.G. performed research; M.Y., H.D., A.R., P.Q. and H.G. analyzed data; and P.Q. and H.G. wrote the paper.

## Competing Interest Statement

The authors declare no conflict of interest.

## Classification

Biological Sciences/Biochemistry.

## Acknowledgments

We thank the late Professor Martin Karplus for a gift of the CHARMM program. This work was partly supported by the Natural Science Foundation of China (No. 22177064 to P.Q.) and the Natural Science Foundation of Shandong Province (No. ZR2021MB050 to P.Q.).

## Notes

### Competing Interest Statement

The authors have declared no competing interest.

### Summary of Updates

1. Title has been changed because the results are more general than previous realized. Reactive conformation used in this version is more commonly used than active induced fit conformation used in the earlier version for better communication with readers. 2. A new author is added who added new data to this new version. 3. The manuscript has been modified significantly to reflect new understanding of the significance of the results. The results/data are basically the same as before with some new simulations data for further confirmation of the conclusions.

