## Supplemental Figure S1, S2 and S3 for "The ability of enzymes to preserve reactive conformations with well-positioned residues during enzyme-catalyzed reactions can be an important factor for efficient catalysis: a case study of CARNMT1"

**This PDF file includes:**

Figures S1 to S3

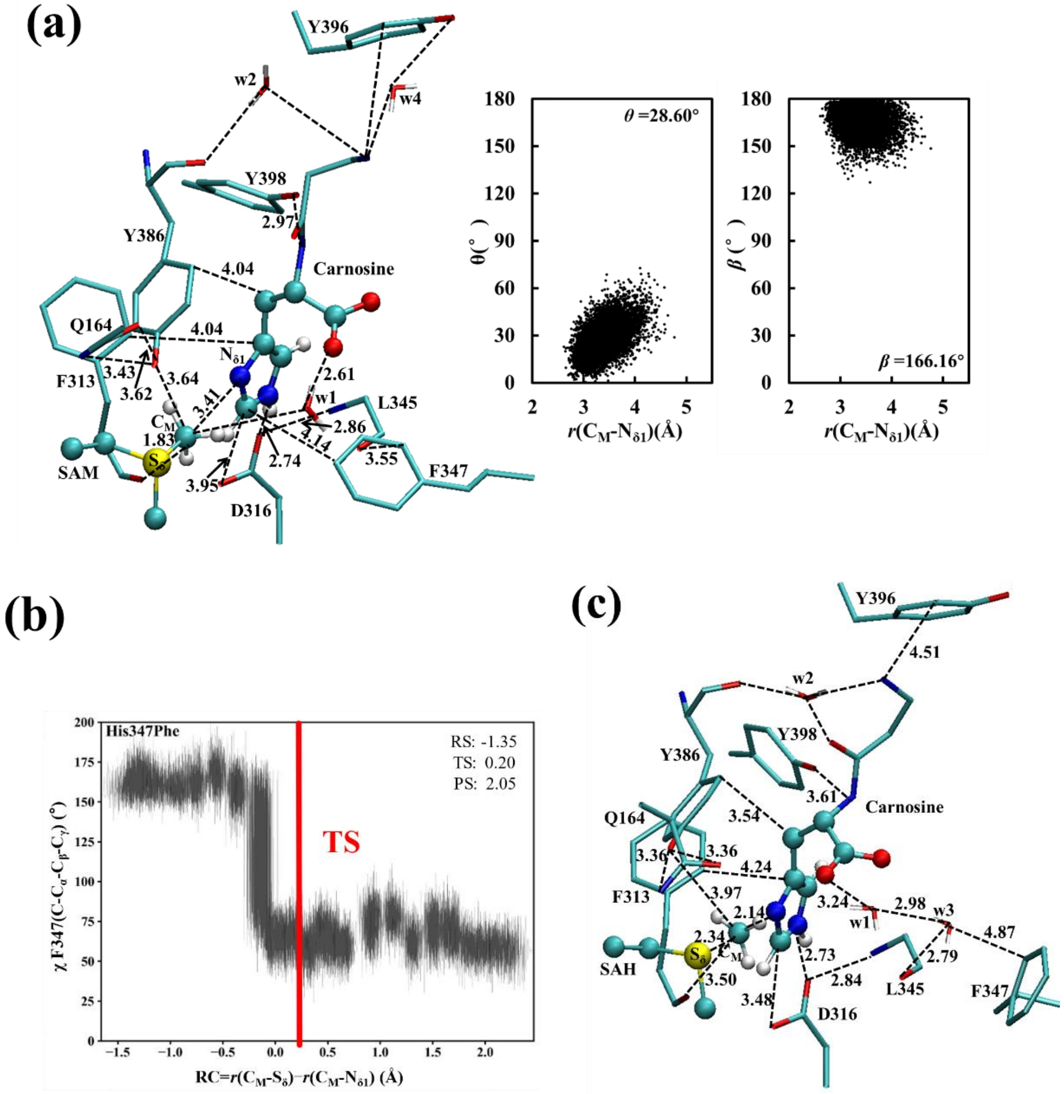


**Fig. S1.** (a) The average structures for the active sites of the His347Phe-substrate complex along with the corresponding distribution maps. (b) The fluctuations of the $\chi(C\text{-}C_\alpha\text{-}C_\beta\text{-}C_\gamma)$ dihedral angle of Phe347 in the PMF windows along the reaction coordinate. (c) The average structure for the active sites near transition state of methylation for the His347Phe.

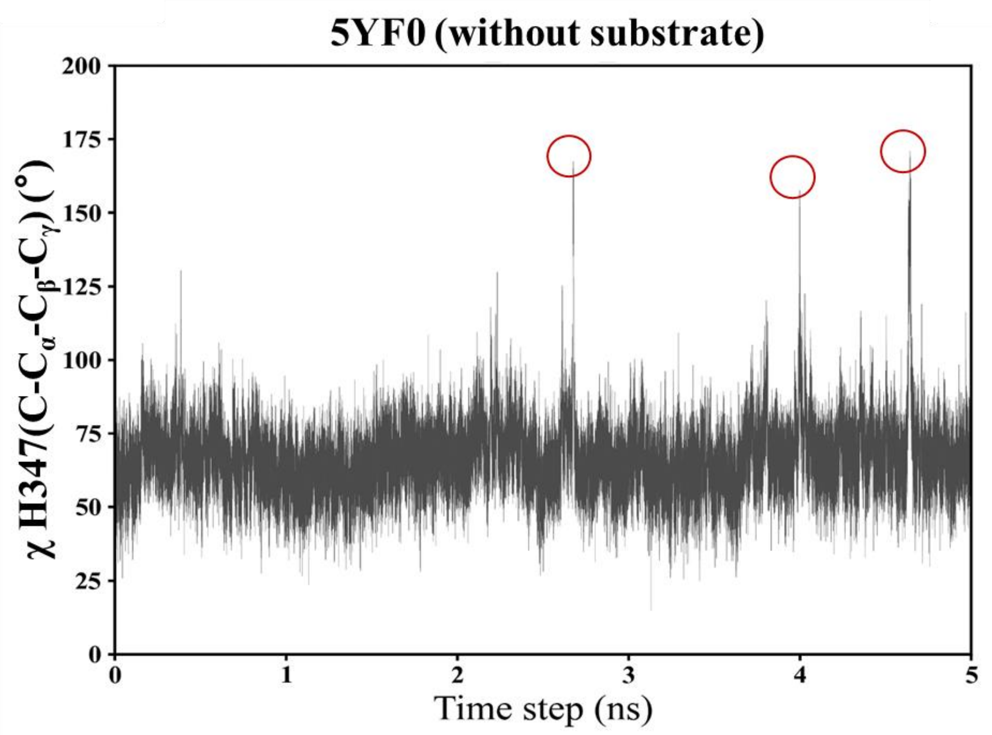


**Fig. S2.** To examine whether the active site residues in the apo enzyme may reach the configurations observed in the reactive conformation of the enzyme-substrate complex, as suggested from the conformational selection model, we performed the QM/MM MD simulations based on 5YF0. The results here show that the residues may reach such configurations as monitored by $\chi(C\text{-}C_{\alpha}\text{-}C_{\beta}\text{-}C_{\gamma})$ (red circles), even though the frequency of formation of such structure without the substrate is quite low.

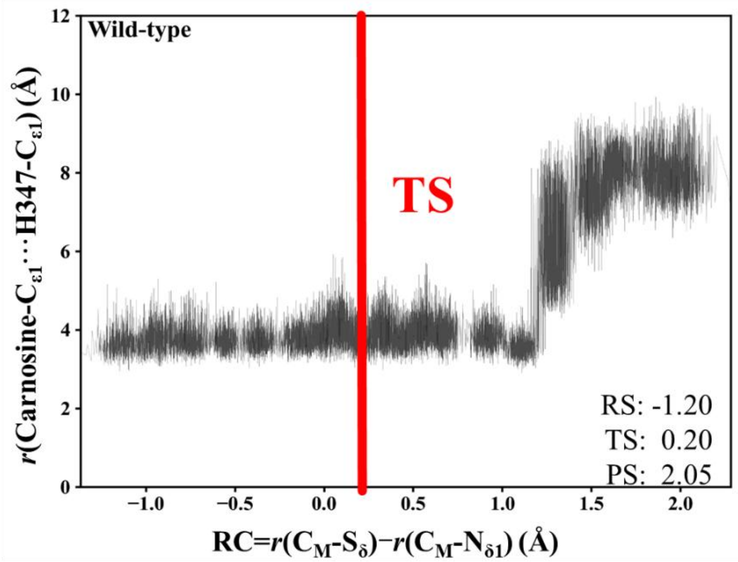


**Figure S3.** The fluctuations of the r(Carnosine-$C_{\varepsilon 1}$···H347-$C_{\varepsilon 1}$) distance in the PMF windows along the reaction coordinate (see Figure 2b).
